# WAC loss alters food-associated behavior and physiological homeostasis with stage-associated cholinergic dysregulation in *Caenorhabditis elegans*

**DOI:** 10.64898/2026.04.17.719318

**Authors:** Da-Woon Kim, Napissara Boonpraman, Nathan C Kuhn, Shreesh Raj Sammi

## Abstract

WAC is a chromatin-associated regulatory protein involved in transcriptional control and has been identified as an autism-associated gene in human genetic studies. However, its functional role in regulating behavior and synaptic processes remains incompletely understood. Using *Caenorhabditis elegans,* we investigated the consequences of *wac* deficiency on food-associated social behavior, growth-associated phenotypes, and cholinergic pathway function. *wac*-deficient worms showed a marked reduction in food-leaving behavior, supporting impaired behavioral responsiveness to food-associated environmental cues, while aggregation behavior was not significantly altered. PHX2587 *wac* deletion mutant worms also exhibited reduced body length, decreased pharyngeal pumping, and shortened lifespan, indicating broader growth and physiological impairment. Stage-resolved analysis of cholinergic pathway genes revealed stage-associated transcriptional changes, with coordinated upregulation of multiple presynaptic and postsynaptic cholinergic components (*ace-1*, *cha-1*, *cho-1*, *lev-1*, *lev-10*, *unc-17*, *unc-29*, *unc-38*, and *unc-50*) emerging most prominently at the young adult stage. Functional RNAi analysis further identified *cho-1*, which encodes the high-affinity presynaptic choline transporter, as a genotype-specific modifier of cholinergic sensitivity in PHX2587 worms. Importantly, *cho-1* RNAi not only reduced aldicarb hypersensitivity but also partially suppressed the reduced body length phenotype and improved food-leaving behavior in PHX2587 worms, while having limited effects in wild-type N2. Together, these findings support a functional relationship between *wac* deficiency and *cho-1*-associated cholinergic modulation, suggesting that presynaptic choline transport contributes to selected behavioral and physiological consequences of *wac* loss.

Graphical Abstract

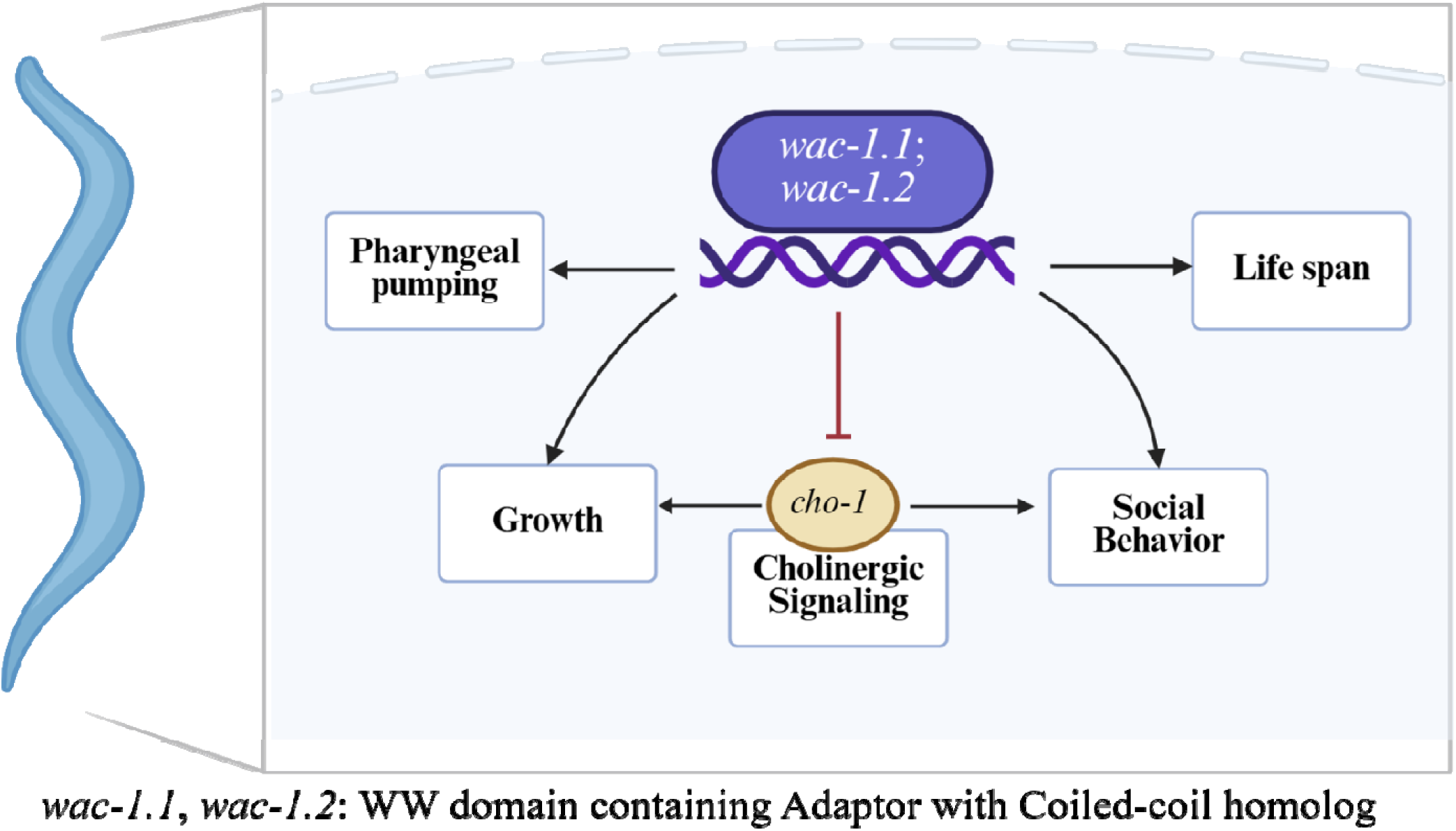

## 1. Introduction

Autism spectrum disorder (ASD) is a neurodevelopmental condition characterized by alterations in social communication and repetitive behavioral patterns, often arising from complex genetic and neurobiological mechanisms (Christensen et al., 2019; Lord et al., 2018). Patients diagnosed with neurodevelopmental disorders, such as ASD, can exhibit altered social behavior and atypical responses to sensory and environmental stimuli (Frye, 2018; Posar and Visconti, 2018; Roley et al., 2015; Thye et al., 2018). Although large-scale genetic studies have identified many ASD-associated genes (Garcia-Forn et al., 2020), an ongoing challenge is to determine how individual risk genes alter specific behavioral domains and to connect those effects to conserved synaptic pathways that translate molecular disruption into circuit-level dysfunction (Bonsi et al., 2022; Sahin and Sur, 2015). WAC (WW domain–containing adaptor with coiled-coil) has been reported as an ASD-associated gene in human genetic studies (Sanders et al., 2015; Satterstrom et al., 2020). Pathogenic variants in WAC are also associated with DeSanto-Shinawi syndrome (DESSH), a neurodevelopmental disorder characterized by intellectual disability and behavioral abnormalities, including autism-related features (Alsahlawi et al., 2020; Leonardi et al., 2020; Lugtenberg et al., 2016; Morales et al., 2022; Uehara et al., 2018). However, reduced WAC function has been associated with diverse behavioral abnormalities, including anxiety, attention-deficit/hyperactivity disorder (ADHD), sleep disturbances, aggression, and autism-related features(Varvagiannis et al., 1993), yet it remains unclear which synaptic mechanisms mediate these effects. At the molecular level, WAC has been linked to key regulatory processes, including chromatin-dependent transcription, mTOR signaling, and autophagy (David-Morrison et al., 2016; Joachim et al., 2015; Zhang and Yu, 2011). More recently, structure–function studies in neuronal systems have further clarified WAC subcellular localization and identified regulatory motifs with potential relevance to neuronal signaling (Rudolph et al., 2023). Nevertheless, translating these molecular insights into testable, mechanism-driven hypotheses linking synaptic dysfunction to behavioral phenotypes remains challenging, highlighting the need for *in vivo* models that pair quantitative behavioral readouts with rigorous mechanistic validation.

*Caenorhabditis elegans* exhibits a broad range of foraging behaviors (Macosko et al., 2009a), and hence provides a tractable *in vivo* model to connect gene function to synaptic mechanisms and behavior (Mizumoto et al., 2023; Rand, 2007). Its nervous system is well characterized, key neurotransmitter pathways are conserved, and behavioral assays can be quantified under tightly controlled developmental staging (Cook et al., 2019b; White et al., 1986a). Food leaving is a widely used assay to quantify food-associated social behavior in *C. elegans* and has been applied to investigate genes linked to neurodevelopmental disorders, including ASD– associated genes (Bendesky et al., 2011; Milward et al., 2011; Rawsthorne et al., 2021b). Previous studies have shown that mutations in ASD-related genes, such as neuroligin (*nlg-1*), alter food-leaving behavior, supporting its utility as a behavioral readout relevant to social and sensory processing (Rawsthorne et al., 2021a; Rawsthorne et al., 2021b). In the present study, VC228/*nlg-1* was included as an assay reference strain for altered food-leaving behavior rather than as a mechanistic comparator for WAC-dependent cholinergic regulation. In parallel, cholinergic signaling can be interrogated using complementary molecular and functional approaches (Mahoney et al., 2006; Rand, 2007). Transcriptional profiling can identify candidate pathway components, and pharmacological assays such as aldicarb sensitivity provide a well-established functional readout of acetylcholine neurotransmission at the neuromuscular junction (Sammi et al., 2022; Sammi et al., 2017).

In our prior study (Boonpraman et al., 2026), we combined behavioral phenotyping in *C. elegans* with complementary analyses in mice and found that WAC perturbation is associated with measurable neurobehavioral changes. In WAC-heterozygous mice, increased cortical expression of CHRNA7 further suggested a potential link between WAC function and cholinergic regulation (Boonpraman et al., 2026). These findings motivated further investigation in the genetically tractable *C. elegans* model to determine whether alterations in cholinergic gene expression translate into measurable functional and behavioral outcomes. Here, we tested the hypothesis that loss of *wac* alters food-associated behavioral responses and cholinergic pathway in *C. elegans*. To address this, we first established baseline developmental and physiological phenotypes to account for potential confounding effects on behavior. We then quantified food-leaving behavior as a cue– dependent food-associated behavioral readout and aggregation as a complementary group-behavior assay. Extending beyond our prior work, which focused primarily on the young adult (YA) stage, we performed stage– resolved transcriptional profiling across larval development (L1–L4) through YA to determine when cholinergic pathway divergence emerges after *wac* loss, consistent with the early-onset nature of neurodevelopmental disorders. Finally, to assess functional relevance, we targeted a subset of upregulated cholinergic genes using RNAi and evaluated their effects on synaptic function using the aldicarb assay, with subsequent focus on *cho-1* as a genotype-specific functional modifier of selected *wac*-associated phenotypes.

## 2. Materials and Methods

### 2.1. Strains, bacterial food, and general culture conditions

Wild-type *C. elegans* (N2), the *wac*-deficient mutant PHX2587 (*wac-1.1*; *wac-1.2*), and VC228 (*nlg-1*) were obtained from the Caenorhabditis Genetics Center (CGC, University of Minnesota). VC228 was included as a positive-control strain for the food-leaving assay(Rawsthorne et al., 2021a; Rawsthorne et al., 2021b). Worms were maintained on standard nematode growth medium (NGM) plates seeded with *Escherichia coli* OP50 and cultured at 22°C. Synchronized cohorts were generated by alkaline hypochlorite treatment of gravid adults to isolate embryos (Porta-de-la-Riva et al., 2012). Embryos were incubated overnight in M9 buffer at 15°C to obtain synchronized L1 larvae. For RNAi feeding experiments (Kamath et al., 2001), RNAi clones were grown overnight at 37°C in Luria broth supplemented with carbenicillin (25 µg/mL). Cultures were then seeded onto NGM plates containing IPTG (1 mM) and ampicillin (50 µg/mL) and incubated overnight at 37°C to induce dsRNA expression. Synchronized L1 larvae were transferred onto RNAi plates and grown at 22°C for the indicated duration prior to downstream assays. *E. coli* HT115(DE3) carrying the empty vector L4440 was used as the negative control.

### 2.2. Quantitative real-time PCR (qRT–PCR) and genomic PCR validation

Quantitative real-time PCR was performed largely as described in our prior study (Boonpraman et al., 2026). Briefly, synchronized N2 and PHX2587 worms were collected at the indicated developmental stages, washed to remove bacteria, and processed for total RNA isolation. cDNA was synthesized using oligo(dT) priming, and qPCR was run on a Bio-Rad CFX96 instrument using a SYBR-based detection chemistry. Each reaction contained 150 ng cDNA. Relative expression was calculated using the 2^−ΔΔCt^ method with *gpd-1* as the internal reference (Livak and Schmittgen, 2001). For developmental comparisons, N2 at each corresponding stage was used as the calibrator (set to 1). Primer sequences were taken from our prior study (Boonpraman et al., 2026). For each biological replicate, approximately 5,000 worms per plate were collected and processed. Experiments were performed with n = 3 independent biological replicates (plates), and each sample was analyzed in technical duplicates by qRT-PCR. To validate the PHX2587 *wac* deletion region, genomic PCR was performed using genomic DNA isolated from N2 and PHX2587 worms. WT-specific primer pairs were designed to amplify three regions across the corresponding wild-type *wac* genomic locus, including the 5′ region, an internal region, and the 3′ region. Primer sequences used for genomic PCR validation are listed in Supplementary Table 1.

### 2.3. Body length measurements and representative imaging

Synchronized L1 larvae were grown on OP50-seeded NGM plates at 22°C, and body length was quantified at the indicated time points (19–53 h). Worms were imaged using a standard light microscope, and body length was measured in ImageJ (v1.54, NIH) using calibrated scale settings. Representative images of N2 and PHX2587 were acquired at matched developmental stages under identical imaging conditions.

### 2.4. Pharyngeal pumping assay

Pharyngeal pumping was measured at 48 h after L1 synchronization (young adult stage). Individual worms were transferred to freshly seeded OP50 NGM plates and pumping was scored while worms were feeding, as described previously(Raizen et al., 2012; Rawsthorne et al., 2021b). Pumps were counted for 20 s under a dissecting microscope and converted to pumps per minute. Measurements were collected from three independent biological replicates (n = 3), with 10 worms scored per group per replicate (total n = 30 worms per group).

### 2.5. Lifespan assay

Lifespan was measured using synchronized populations cultured at 22°C, following established procedures with minor modifications (Solis and Petrascheck, 2011). Day 0 was defined as the day synchronized L1 larvae were transferred onto OP50-seeded NGM plates. Worms were monitored until all worms were dead, up to day 25. Beginning at the young adult stage, worms were transferred to fresh OP50-seeded NGM plates daily to prevent confounding by progeny. Survival was scored at regular intervals, and worms were censored if lost or if they died from non-aging causes. Kaplan–Meier survival curves were generated and compared using the log-rank (Mantel–Cox) test. Survival data were compiled from three independent biological replicates, with a total of 261 N2 worms and 313 PHX2587 worms analyzed.

### 2.6. Food-leaving behavior assay

Food-leaving behavior was quantified using a modified version of the assay described by Rawsthorne et al. (2021). Briefly, OP50 lawns of uniform size and bacterial density were pre-conditioned with 100–200 N2 progeny prior to the assay. To generate progeny-conditioned lawns, synchronized gravid N2 adults were placed onto OP50 lawns and allowed to lay progeny for a defined period, after which the adults were removed. Synchronized young adult worms of each genotype (N2, PHX2587, and VC228) were then transferred onto the progeny-conditioned lawns and allowed to acclimate. Food-leaving behavior was scored at defined time points after placement on food (2 h and 24 h). For RNAi-based food-leaving assays, synchronized L1 worms were grown on empty vector (EV), *wac-1.2*, or *cho-1* RNAi plates until the young adult stage. Stage-matched young adult worms were then transferred to OP50 assay lawns containing N2 progeny. Leaving was defined as a worm that had completely moved off the bacterial lawn at the time of scoring. Data are presented as the proportion of worms off the lawn per plate, averaged across plates within each independent biological replicate. For each independent biological replicate (n = 3 plates), 10 worms per genotype were analyzed in duplicate technical replicates, and the proportion of worms off the lawn was calculated at each scoring time point.

### 2.7. Aggregation assay

Aggregation on OP50 lawns was quantified using a previously described protocol with minor modifications (Cowen et al., 2024). Briefly, L1-synchronized worms were cultured on OP50-seeded NGM plates at 22°C and scored at 48 h after synchronization, corresponding to the young adult stage. Aggregation was defined as a cluster of three or more worms in direct body contact. Because individual worms within dense aggregates could not always be reliably distinguished, each aggregate cluster was counted as one aggregation event. The total number of worms present on each plate was counted, and aggregation was expressed as a normalized aggregation index by dividing the number of aggregation events by the total number of worms on the plate. For each biological replicate, 100–150 worms were scored per strain. Three independent biological replicates were performed. Data are presented as mean ± SEM. Statistical analysis was performed using ordinary one-way ANOVA followed by Tukey’s multiple-comparisons test.

### 2.8. Aldicarb sensitivity assay

Cholinergic synaptic function was evaluated using an aldicarb-induced paralysis assay performed largely as described previously (Sammi et al., 2022). Age-synchronized young adults were washed and transferred to NGM plates supplemented with 0.5 mM aldicarb. Aldicarb inhibits acetylcholinesterase, thereby increasing acetylcholine levels at the neuromuscular junction and producing a time-dependent paralysis phenotype. Accordingly, the kinetics of aldicarb-induced paralysis provide a widely used functional readout of cholinergic neurotransmission in *C. elegans* (Mahoney et al., 2006). Paralysis was scored at regular intervals (e.g., every 30 min) using a standardized criterion (lack of movement in response to gentle prodding). Worms lost or injured during handling were excluded from analysis. Experiments were performed with six independent biological replicates, each containing 30-40 worms.

### 2.9. Statistical Analysis

Statistical analyses were performed using GraphPad Prism 10 (GraphPad Software, La Jolla, CA, USA). Experiments were conducted with at least three independent biological replicates unless otherwise stated, and data are presented as mean ± SEM. Statistical comparisons were performed using the appropriate test for each experiment, including unpaired t-tests, one-way or two-way ANOVA followed by multiple-comparisons testing, or log-rank (Mantel–Cox) analysis for survival data, as indicated in the figure legends. A p-value < 0.05 was considered statistically significant.

### 2.10. The C. elegans Neuronal Gene Expression Map & Network (CeNGEN) analysis of wac-1.1 and wac-1.2 in cholinergic neurons

Public CeNGEN single-cell RNA-seq data were used to examine the expression profiles of *wac-1.1* and *wac-1.2* across cholinergic neuron classes in *C. elegans*. Gene expression values were queried using the CeNGEN(Taylor et al., 2021). Query was run using the gene sequence names: Y40B1A.3 for *wac-1.1* and Y40B1A.1 for *wac-1.2*. Expression was examined across the available L1, L4, and adult hermaphrodite neuronal datasets. Cholinergic neuron classes were selected based on published acetylcholine neuron annotations(Hammarlund et al., 2018). For each gene, two values per cell type were recorded: the mean expression across all cells of that type, in transcripts per million (TPM), and the fraction of cells in which the transcript was detected, reported as a percentage. Detection was called at a threshold of 2, which is the default on 1–4 stringency scale of CENGEN(Taylor et al., 2021). Tables were generated for every annotated cell type at each stage, which were then narrowed to the neurons of interest. Any non– neuronal cell types in the L4 data were dropped from the analysis. Heatmaps comparing the two genes, wac-1.1 and wac-1.2 across cholinergic classes and stages in python 3 (pandas, Matplotlib), plotting the TPM and expressing percent metric on separate panels.

## 3. Results

### Loss of wac reduces food-leaving behavior while aggregation behavior remains unaltered

Before behavioral and functional analyses, we validated the PHX2587 *wac* deletion region by genomic PCR using WT-specific primer sets spanning the corresponding 5′, internal, and 3′ regions of the wild-type *wac* locus. Expected PCR products were detected in N2 genomic DNA but not in PHX2587 genomic DNA, consistent with loss of the corresponding WT *wac* genomic regions in PHX2587 worms (**Supplementary Fig. 1**). To determine whether loss of *wac* affects food-associated social behaviors, we quantified food-leaving behavior on OP50 assay lawns containing N2 progeny. Compared with wild-type N2, *wac*-deficient PHX2587 worms exhibited reduced food leaving at 24 h after placement on progeny-containing OP50 lawns (N2: 48.72 ± 5.13%; PHX2587: 9.44 ± 0.55%) (**Fig. 1A**). In N2 worms, food leaving increased over time (2 h: 7.69 ± 0.00% to 24 h: 48.72 ± 5.13%), whereas PHX2587 showed minimal change (2 h: 0.00 ± 0.00% to 24 h: 9.44 ± 0.55%), indicating a blunted temporal response in the mutant strain. The VC228/*nlg-1* assay reference strain displayed the expected reduced food-leaving phenotype (2 h: 0.00 ± 0.00% to 24 h: 10.00 ± 5.77%), consistent with prior reports (Rawsthorne et al., 2021a; Rawsthorne et al., 2021b). Notably, the magnitude of reduction observed in PHX2587 worms was comparable to that of VC228, further supporting a robust defect in food-leaving behavior upon *wac* loss. We next quantified aggregation behavior as a complementary group-behavior assay. Unlike food-leaving behavior, aggregation behavior was not significantly altered among N2, PHX2587, and VC228 worms under these assay conditions (**Fig. 1B**). These findings indicate that loss of *wac* selectively reduces food– leaving behavior without producing a detectable change in aggregation behavior in this assay.

**Figure 1.**
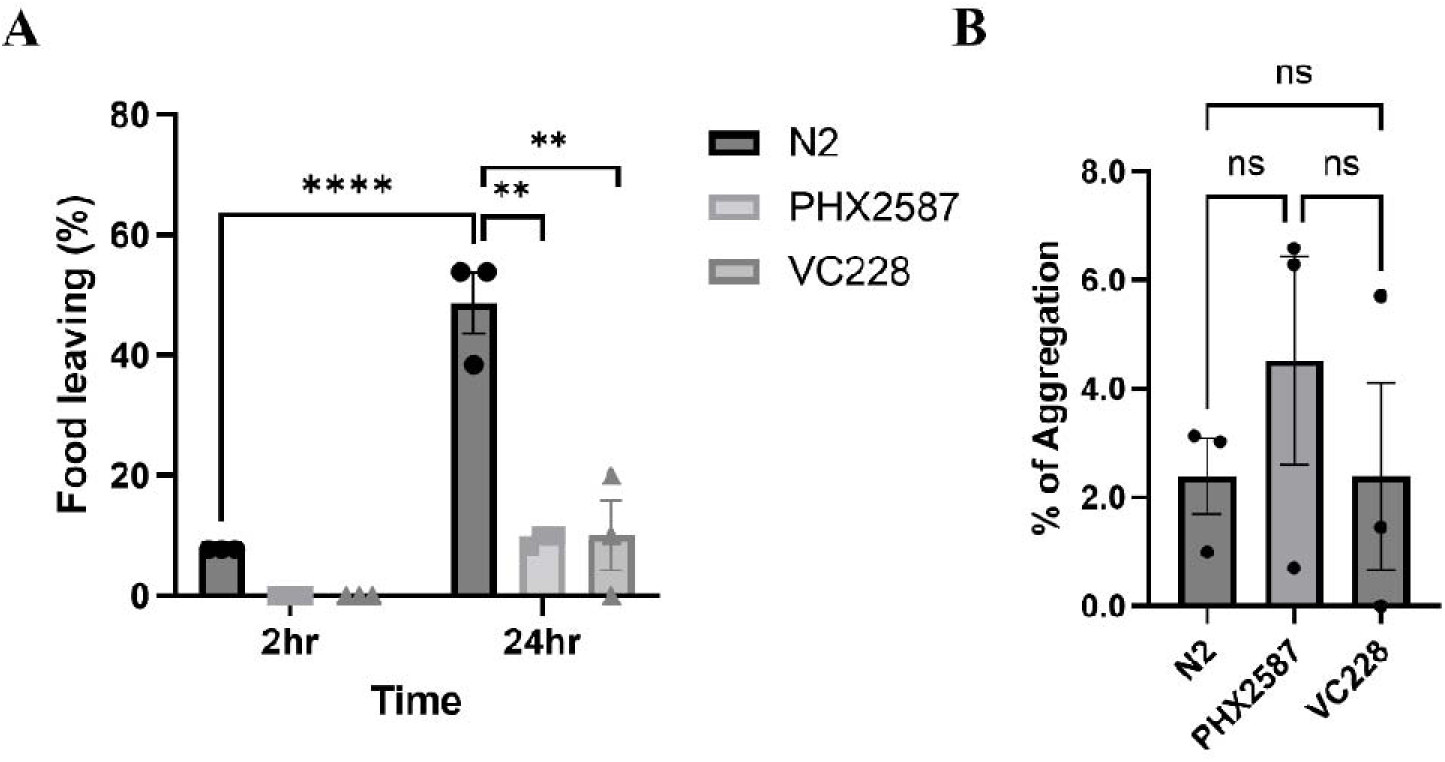
Analysis of food-leaving and aggregation behavior following loss of wac in C. elegans. (A) Food– leaving behavior of wild-type N2, *wac*-deficient PHX2587, and the *nlg-1* deficient VC228 assay reference strain was measured at 2 h and 24 h after placement on standardized OP50 lawns containing N2 progeny. Each dot represents an independent biological replicate (n = 3 plates), with 10 worms scored per genotype in each replicate. Bars indicate mean ± SEM. Data were analyzed by two-way ANOVA followed by Šidák’s multiple– comparisons test. Statistical significance is indicated as **p < 0.01 and ****p < 0.0001. (B) Aggregation behavior of N2, PHX2587, and VC228 worms was quantified at the young adult stage under identical assay conditions. Aggregation was defined as a cluster of three or more worms in direct body contact, and each aggregate cluster was counted as one aggregation event. Each dot represents an independent biological replicate (n = 3 plates), with 100–150 worms scored per replicate. Bars indicate mean ± SEM. Data were analyzed by ordinary one-way ANOVA followed by Tukey’s multiple-comparisons test. ns, not significant.

### Loss of wac is associated with stage-associated changes in cholinergic pathway gene expression

To further assess whether *wac* transcripts are detectable in cholinergic neuronal populations, we examined public CeNGEN single-cell RNA-seq datasets. Using the corresponding WormBase sequence names, *wac-1.1* and *wac-1.2* transcripts were detected across multiple cholinergic neuron classes at L1, L4, and adult stages (**Supplementary Fig. 2**). These public expression data support the plausibility that *wac* function may intersect with cholinergic circuits, although they do not establish cell-autonomous function or direct regulation of cholinergic pathway genes by WAC. To determine whether loss of *wac* is associated with transcriptional changes relevant to cholinergic signaling, we quantified expression of a predefined panel of cholinergic pathway genes (Boonpraman et al., 2026) across larval stages (L1–L4) and young adults (YA) in synchronized N2 and PHX2587 worms (**Fig. 2**; **Table 1**). A global view of transcriptional changes is presented in the heatmap (**Fig. 2A**), which displays log2(PHX2587/N2) values across development. Consistent with our prior study (Boonpraman et al., 2026), in which coordinated changes were primarily detected at the YA stage, PHX2587 exhibited a pronounced late-stage shift, with the majority of cholinergic pathway genes showing increased expression relative to N2. Gene-level resolution (**Fig. 2B**) revealed that at the YA stage, 9 of the 14 genes were significantly altered relative to N2, all of which were upregulated. These included *ace-1* (fold change [FC] = 1.97), *cha-1* (FC = 1.91), *cho-1* (FC = 1.98), *lev-1* (FC = 8.25), *lev-10* (FC = 5.34), *unc-17* (FC = 2.07), *unc-29* (FC = 2.90), *unc-38* (FC = 5.23), and *unc-50* (FC = 2.98) (Fig. 2B; Table 1). Notably, several genes, particularly *lev-1*, *lev-10*, and *unc-38*, exhibited strong upregulation at YA stage, indicating coordinated upregulation of both pre– and postsynaptic components of cholinergic signaling. Importantly, extending the analysis across larval development revealed additional stage-associated patterns that are not apparent from YA– only analysis. Select genes showed early divergence prior to adulthood: *ace-1* and *acr-3* were upregulated at L1, no genes were significantly altered at L2, *acr-3* was upregulated at L3, and *cha-1*, *cho-1*, *lev-1*, and *lev-10* were upregulated at L4 (**Fig. 2B**; **Table 1**). Together, these data indicate that while the most pronounced and coordinated transcriptional changes occur at the YA stage, loss of *wac* is associated with earlier gene– and stage-associated divergence within the cholinergic pathway.

**Figure 2.**
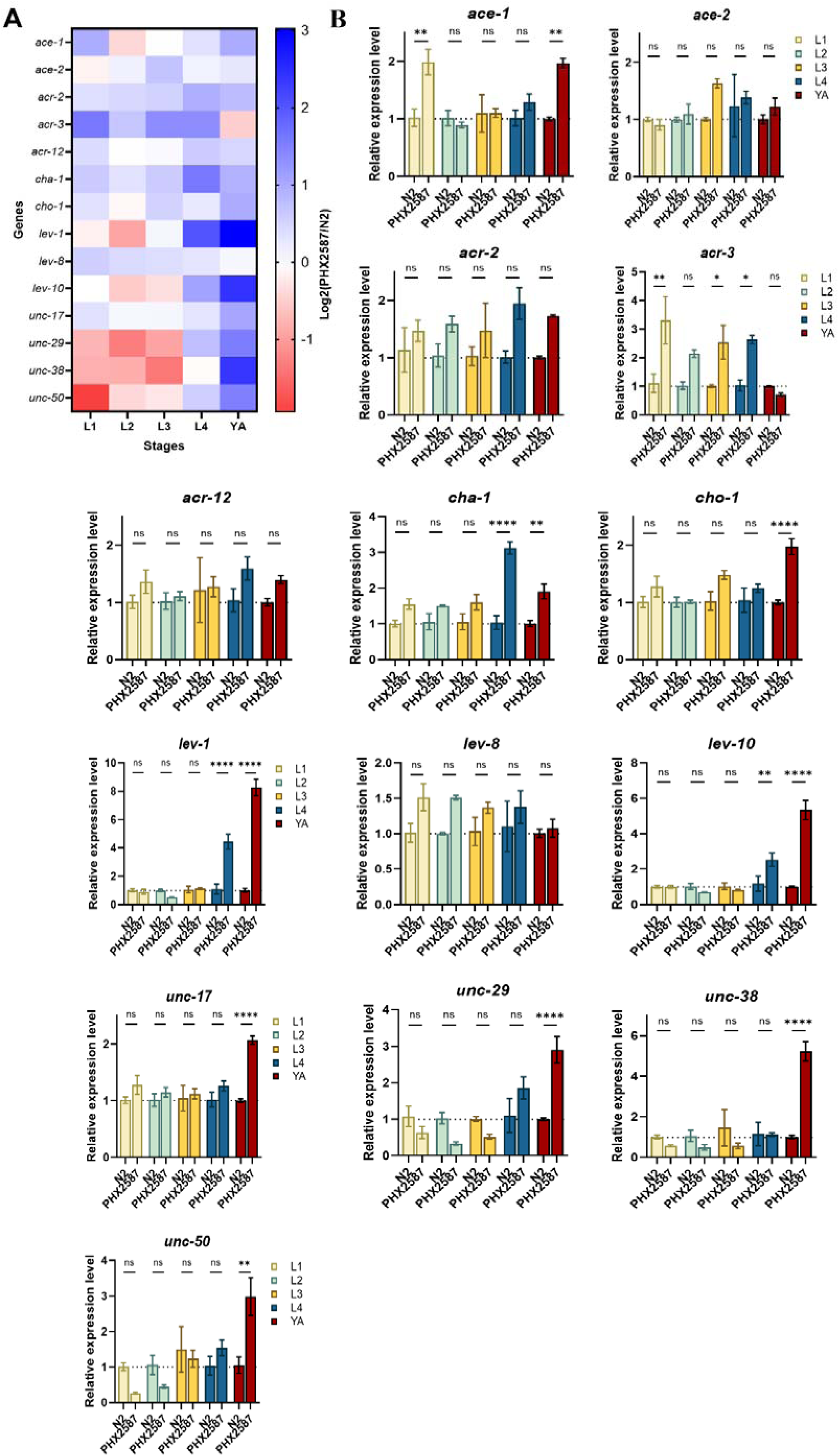
Stage-resolved analysis of cholinergic pathway gene expression in wac-deficient C. elegans. (A) Heatmap showing stage-resolved relative expression changes in cholinergic pathway genes across larval stages (L1–L4) and young adults (YA), displayed as log2(PHX2587/N2) values derived from qRT–PCR measurements. Each row represents a gene and each column represents a developmental stage, allowing visualization of stage-associated expression dynamics across the developmental timeline. (B) Relative mRNA expression levels of individual cholinergic pathway genes in N2 and PHX2587 across developmental stages (L1–L4 and YA). For panel B, expression values were normalized to N2 at each corresponding developmental stage, with N2 set to 1 at each stage (dashed line = 1), to enable stage-matched genotype comparisons between N2 and PHX2587. Data are presented as mean ± SEM from three independent biological replicates (plates), with each sample analyzed in technical duplicates by qRT–PCR. Statistical analysis was performed using two– way ANOVA (genotype × developmental stage), followed by Šidák’s multiple comparisons test to compare N2 and PHX2587 at each stage. Statistical significance is indicated as *p < 0.05, **p < 0.01, and ****p < 0.0001.

**Table 1.**
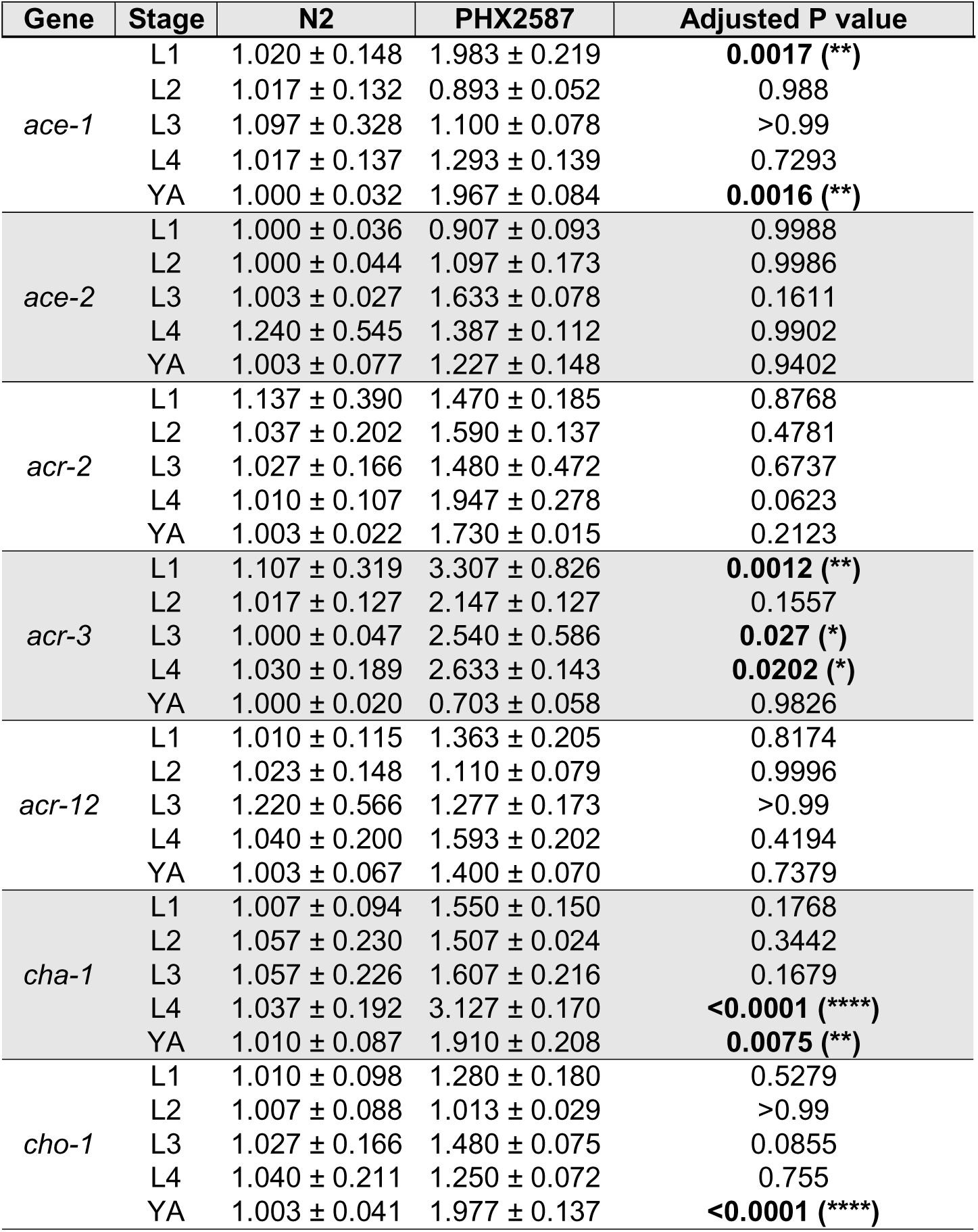
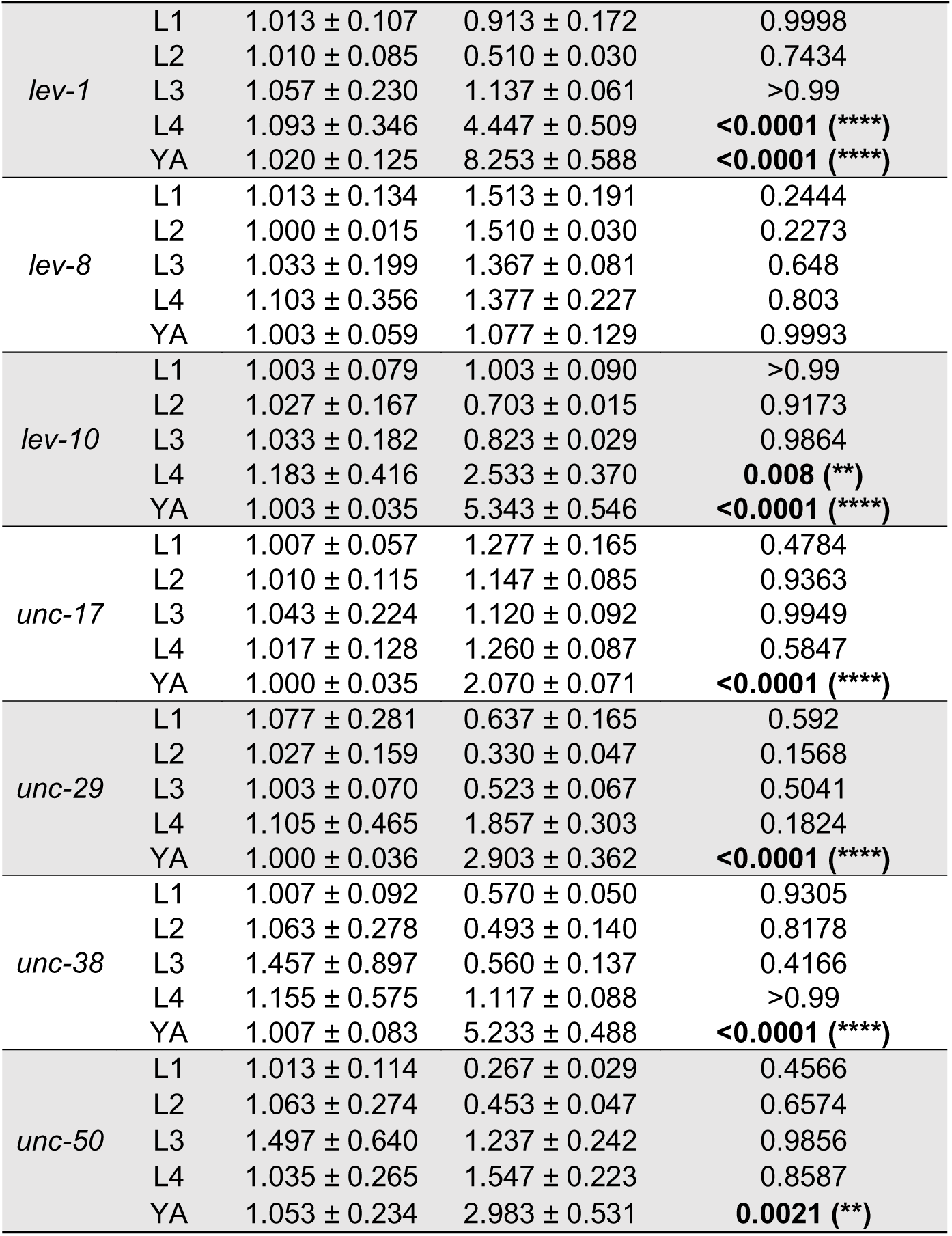
Stage-resolved qRT–PCR analysis of cholinergic pathway gene expression in wac-deficient C. elegans. Relative mRNA expression levels of cholinergic pathway genes in N2 and PHX2587 worms across developmental stages (L1–L4 and young adult, YA). Expression values were normalized to N2 at the corresponding developmental stage and calculated using the 2^−ΔΔCt method with *gpd-1* as the internal reference. Data are presented as mean relative expression ± SEM from three independent biological replicates (plates), with approximately 5,000 worms collected per replicate and each sample analyzed in technical duplicates by qRT–PCR. Adjusted P values were calculated using two-way ANOVA (genotype × developmental stage) followed by Šidák’s multiple comparisons test to compare N2 and PHX2587 at each developmental stage. Significant values (P < 0.05) are highlighted in bold.

| Gene | Stage | N2 | PHX2587 | Adjusted P value |
| --- | --- | --- | --- | --- |
| *ace-1* | L1 | 1.020 ± 0.148 | 1.983 ± 0.219 | **0.0017 (\*\*)** |
| | L2 | 1.017 ± 0.132 | 0.893 ± 0.052 | 0.988 |
| | L3 | 1.097 ± 0.328 | 1.100 ± 0.078 | >0.99 |
| | L4 | 1.017 ± 0.137 | 1.293 ± 0.139 | 0.7293 |
| | YA | 1.000 ± 0.032 | 1.967 ± 0.084 | **0.0016 (\*\*)** |
| *ace-2* | L1 | 1.000 ± 0.036 | 0.907 ± 0.093 | 0.9988 |
| | L2 | 1.000 ± 0.044 | 1.097 ± 0.173 | 0.9986 |
| | L3 | 1.003 ± 0.027 | 1.633 ± 0.078 | 0.1611 |
| | L4 | 1.240 ± 0.545 | 1.387 ± 0.112 | 0.9902 |
| | YA | 1.003 ± 0.077 | 1.227 ± 0.148 | 0.9402 |
| *acr-2* | L1 | 1.137 ± 0.390 | 1.470 ± 0.185 | 0.8768 |
| | L2 | 1.037 ± 0.202 | 1.590 ± 0.137 | 0.4781 |
| | L3 | 1.027 ± 0.166 | 1.480 ± 0.472 | 0.6737 |
| | L4 | 1.010 ± 0.107 | 1.947 ± 0.278 | 0.0623 |
| | YA | 1.003 ± 0.022 | 1.730 ± 0.015 | 0.2123 |
| *acr-3* | L1 | 1.107 ± 0.319 | 3.307 ± 0.826 | **0.0012 (\*\*)** |
| | L2 | 1.017 ± 0.127 | 2.147 ± 0.127 | 0.1557 |
| | L3 | 1.000 ± 0.047 | 2.540 ± 0.586 | **0.027 (\*)** |
| | L4 | 1.030 ± 0.189 | 2.633 ± 0.143 | **0.0202 (\*)** |
| | YA | 1.000 ± 0.020 | 0.703 ± 0.058 | 0.9826 |
| *acr-12* | L1 | 1.010 ± 0.115 | 1.363 ± 0.205 | 0.8174 |
| | L2 | 1.023 ± 0.148 | 1.110 ± 0.079 | 0.9996 |
| | L3 | 1.220 ± 0.566 | 1.277 ± 0.173 | >0.99 |
| | L4 | 1.040 ± 0.200 | 1.593 ± 0.202 | 0.4194 |
| | YA | 1.003 ± 0.067 | 1.400 ± 0.070 | 0.7379 |
| *cha-1* | L1 | 1.007 ± 0.094 | 1.550 ± 0.150 | 0.1768 |
| | L2 | 1.057 ± 0.230 | 1.507 ± 0.024 | 0.3442 |
| | L3 | 1.057 ± 0.226 | 1.607 ± 0.216 | 0.1679 |
| | L4 | 1.037 ± 0.192 | 3.127 ± 0.170 | **<0.0001 (\*\*\*\*)** |
| | YA | 1.010 ± 0.087 | 1.910 ± 0.208 | **0.0075 (\*\*)** |
| *cho-1* | L1 | 1.010 ± 0.098 | 1.280 ± 0.180 | 0.5279 |
| | L2 | 1.007 ± 0.088 | 1.013 ± 0.029 | >0.99 |
| | L3 | 1.027 ± 0.166 | 1.480 ± 0.075 | 0.0855 |
| | L4 | 1.040 ± 0.211 | 1.250 ± 0.072 | 0.755 |
| | YA | 1.003 ± 0.041 | 1.977 ± 0.137 | **<0.0001 (\*\*\*\*)** |

**Table 1. Continued.**
| Gene | Stage | N2 | PHX2587 | Adjusted P value |
| --- | --- | --- | --- | --- |
| *lev-1* | L1 | 1.013 ± 0.107 | 0.913 ± 0.172 | 0.9998 |
|  | L2 | 1.010 ± 0.085 | 0.510 ± 0.030 | 0.7434 |
|  | L3 | 1.057 ± 0.230 | 1.137 ± 0.061 | >0.99 |
|  | L4 | 1.093 ± 0.346 | 4.447 ± 0.509 | **<0.0001 (\*\*\*\*)** |
|  | YA | 1.020 ± 0.125 | 8.253 ± 0.588 | **<0.0001 (\*\*\*\*)** |
| *lev-8* | L1 | 1.013 ± 0.134 | 1.513 ± 0.191 | 0.2444 |
|  | L2 | 1.000 ± 0.015 | 1.510 ± 0.030 | 0.2273 |
|  | L3 | 1.033 ± 0.199 | 1.367 ± 0.081 | 0.648 |
|  | L4 | 1.103 ± 0.356 | 1.377 ± 0.227 | 0.803 |
|  | YA | 1.003 ± 0.059 | 1.077 ± 0.129 | 0.9993 |
| *lev-10* | L1 | 1.003 ± 0.079 | 1.003 ± 0.090 | >0.99 |
|  | L2 | 1.027 ± 0.167 | 0.703 ± 0.015 | 0.9173 |
|  | L3 | 1.033 ± 0.182 | 0.823 ± 0.029 | 0.9864 |
|  | L4 | 1.183 ± 0.416 | 2.533 ± 0.370 | **0.008 (\*\*)** |
|  | YA | 1.003 ± 0.035 | 5.343 ± 0.546 | **<0.0001 (\*\*\*\*)** |
| *unc-17* | L1 | 1.007 ± 0.057 | 1.277 ± 0.165 | 0.4784 |
|  | L2 | 1.010 ± 0.115 | 1.147 ± 0.085 | 0.9363 |
|  | L3 | 1.043 ± 0.224 | 1.120 ± 0.092 | 0.9949 |
|  | L4 | 1.017 ± 0.128 | 1.260 ± 0.087 | 0.5847 |
|  | YA | 1.000 ± 0.035 | 2.070 ± 0.071 | **<0.0001 (\*\*\*\*)** |
| *unc-29* | L1 | 1.077 ± 0.281 | 0.637 ± 0.165 | 0.592 |
|  | L2 | 1.027 ± 0.159 | 0.330 ± 0.047 | 0.1568 |
|  | L3 | 1.003 ± 0.070 | 0.523 ± 0.067 | 0.5041 |
|  | L4 | 1.105 ± 0.465 | 1.857 ± 0.303 | 0.1824 |
|  | YA | 1.000 ± 0.036 | 2.903 ± 0.362 | **<0.0001 (\*\*\*\*)** |
| *unc-38* | L1 | 1.007 ± 0.092 | 0.570 ± 0.050 | 0.9305 |
|  | L2 | 1.063 ± 0.278 | 0.493 ± 0.140 | 0.8178 |
|  | L3 | 1.457 ± 0.897 | 0.560 ± 0.137 | 0.4166 |
|  | L4 | 1.155 ± 0.575 | 1.117 ± 0.088 | >0.99 |
|  | YA | 1.007 ± 0.083 | 5.233 ± 0.488 | **<0.0001 (\*\*\*\*)** |
| *unc-50* | L1 | 1.013 ± 0.114 | 0.267 ± 0.029 | 0.4566 |
|  | L2 | 1.063 ± 0.274 | 0.453 ± 0.047 | 0.6574 |
|  | L3 | 1.497 ± 0.640 | 1.237 ± 0.242 | 0.9856 |
|  | L4 | 1.035 ± 0.265 | 1.547 ± 0.223 | 0.8587 |
|  | YA | 1.053 ± 0.234 | 2.983 ± 0.531 | **0.0021 (\*\*)** |

### RNAi knockdown of the cholinergic pathway genes identifies cho-1 as a genotype-specific modifier of aldicarb-induced paralysis

To functionally evaluate the cholinergic pathway genes, we performed RNAi feeding followed by aldicarb-induced paralysis assay in N2 and PHX2587 worms. In N2, RNAi targeting *acr-3*, *cha-1*, and *lev-8* significantly reduced paralysis relative to the N2 EV control, whereas *acr-2*, *acr-12*, and *cho-1* did not produce a significant effect (**Fig. 3A**), suggesting a limited contribution to baseline aldicarb sensitivity in wild-type N2 worms. In contrast, in PHX2587, RNAi targeting *acr-2*, *acr-3*, *acr-12*, *cha-1*, and *cho-1* significantly reduced paralysis relative to the PHX2587 EV control, while *lev-8* showed no detectable effect (**Fig. 3B**), suggesting a broader sensitivity of the cholinergic pathway to perturbation in the *wac*-deficient background. Notably, *cho-1* knockdown selectively reduced paralysis in PHX2587 but not in N2 (N2: 35.83 ± 0.30%; PHX2587: 68.80 ± 3.05% relative to EV), identifying *cho-1* as a genotype-specific modifier of aldicarb sensitivity in the *wac*– deficient background. This selective effect further supports a genotype-specific functional role for *cho-1* in PHX2587 worms. Normalization to each genotype’s EV control (RNAi/EV) further highlighted this difference, with *cho-1* RNAi showing reduced paralysis in PHX2587 compared to N2 (N2: 111.10 ± 4.31%; PHX2587: 71.13 ± 3.91%) (**Fig. 3C**), supporting a genotype-dependent functional interaction between *wac* and *cho-1*– associated cholinergic modulation. These RNAi responses are consistent with the gene expression data, in which multiple cholinergic pathway genes, including *cho-1*, showed elevated expression in PHX2587, particularly at the YA stage. Together, these findings identify *cho-1* as a genotype-specific functional modifier of cholinergic sensitivity in PHX2587 worms.

**Figure 3.**
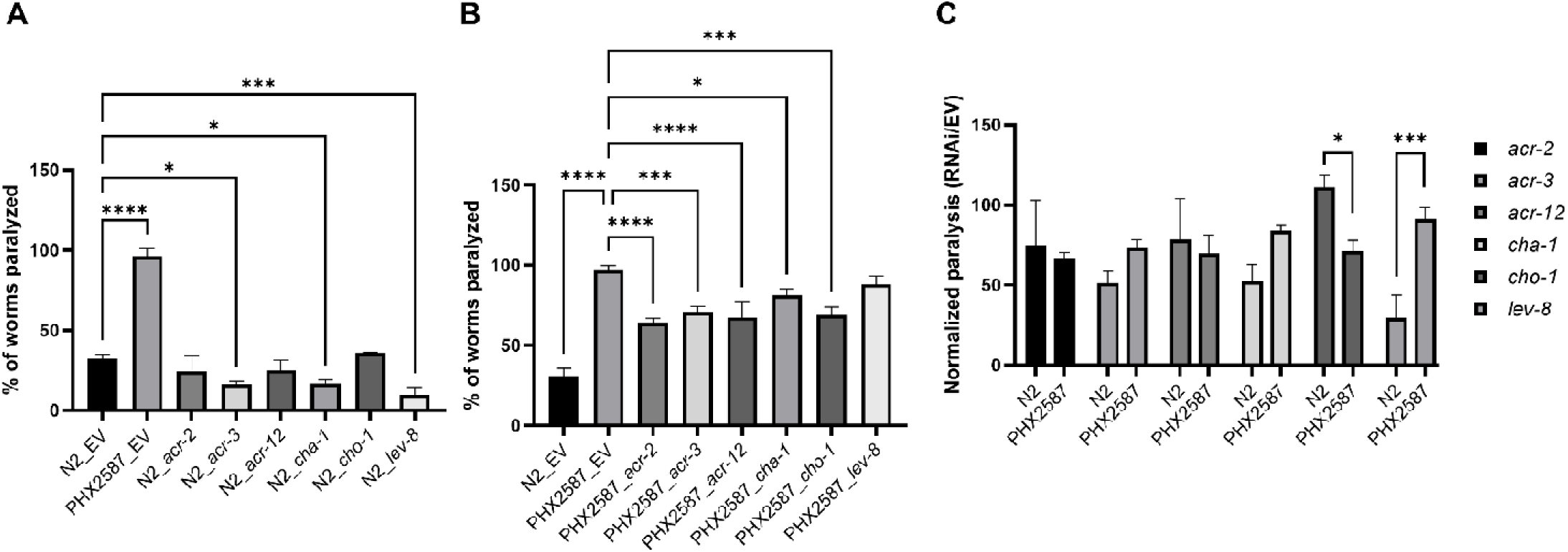
Aldicarb-induced paralysis after RNAi knockdown of cholinergic pathway genes in N2 and wac– deficient PHX2587. (A) Percentage of paralyzed worms in wild-type N2 following RNAi-mediated knockdown of selected cholinergic pathway genes (*acr-2*, *acr-3*, *acr-12*, *cha-1*, *cho-1*, *lev-8*), compared to empty vector (EV) control. Bars represent mean ± SEM from six independent biological replicates, each containing 30–40 worms per condition. Statistical analysis was performed using one-way ANOVA followed by Tukey’s multiple comparisons test. Significant differences compared to N2 EV are indicated as *p < 0.05, ***p < 0.001, and ****p < 0.0001. (B) Percentage of paralyzed worms in *wac*-deficient PHX2587 following RNAi-mediated knockdown of the same gene set, compared to PHX2587 EV control. Bars represent mean ± SEM from six independent biological replicates, each containing 30–40 worms per condition. Statistical analysis was performed using one-way ANOVA followed by Tukey’s multiple comparisons test. Significant differences compared to PHX2587 EV are indicated as *p < 0.05, ***p < 0.001, and ****p < 0.0001. (C) Normalized paralysis ratio calculated as (RNAi / EV). Bars represent mean ± SEM from six independent biological replicates, each containing 30–40 worms per condition. Statistical analysis was performed using one-way ANOVA followed by Tukey’s multiple comparisons test. Significant differences between N2 and PHX2587 are indicated as *p < 0.05 and ***p < 0.001.

### Growth, feeding physiology, and lifespan are impaired by loss of wac in C. elegans

To define baseline growth-associated and physiological phenotypes associated with loss of *wac*, we quantified body length, pharyngeal pumping, and lifespan in wild-type N2 and *wac*-deficient PHX2587 worms, three established readouts of growth, feeding physiology, and organismal fitness in *C. elegans*. After L1 synchronization, body length did not differ at early time points (19 h: N2, ∼523 ± 50 µm; PHX2587, ∼520 ± 70 µm). However, PHX2587 worms were significantly smaller than N2 at later time points, with differences emerging at 43 h (N2: ∼1227 ± 120 µm; PHX2587: ∼1038 ± 120 µm) and persisting through 53 h (N2: ∼1658 ± 130 µm; PHX2587: ∼1375 ± 120 µm) (**Fig. 4A**). Representative images at matched developmental stages were consistent with these body length differences (**Fig. 4B**). To assess feeding-related physiology, we measured pharyngeal pumping at 48 h after L1 synchronization. PHX2587 exhibited a marked reduction in pumping rate relative to N2 (N2: 174.4 ± 18.9 pumps/min; PHX2587: 140.2 ± 16.6 pumps/min) (**Fig. 4C**). Consistent with these findings, lifespan analysis revealed reduced survival in PHX2587 compared with N2, with median survival decreasing from 12 days in N2 (n = 261) to 6 days in PHX2587 (n = 313) (**Fig. 4D**). These results indicate that loss of *wac* is associated with impaired growth-associated progression, feeding physiology, and survival.

**Figure 4.**
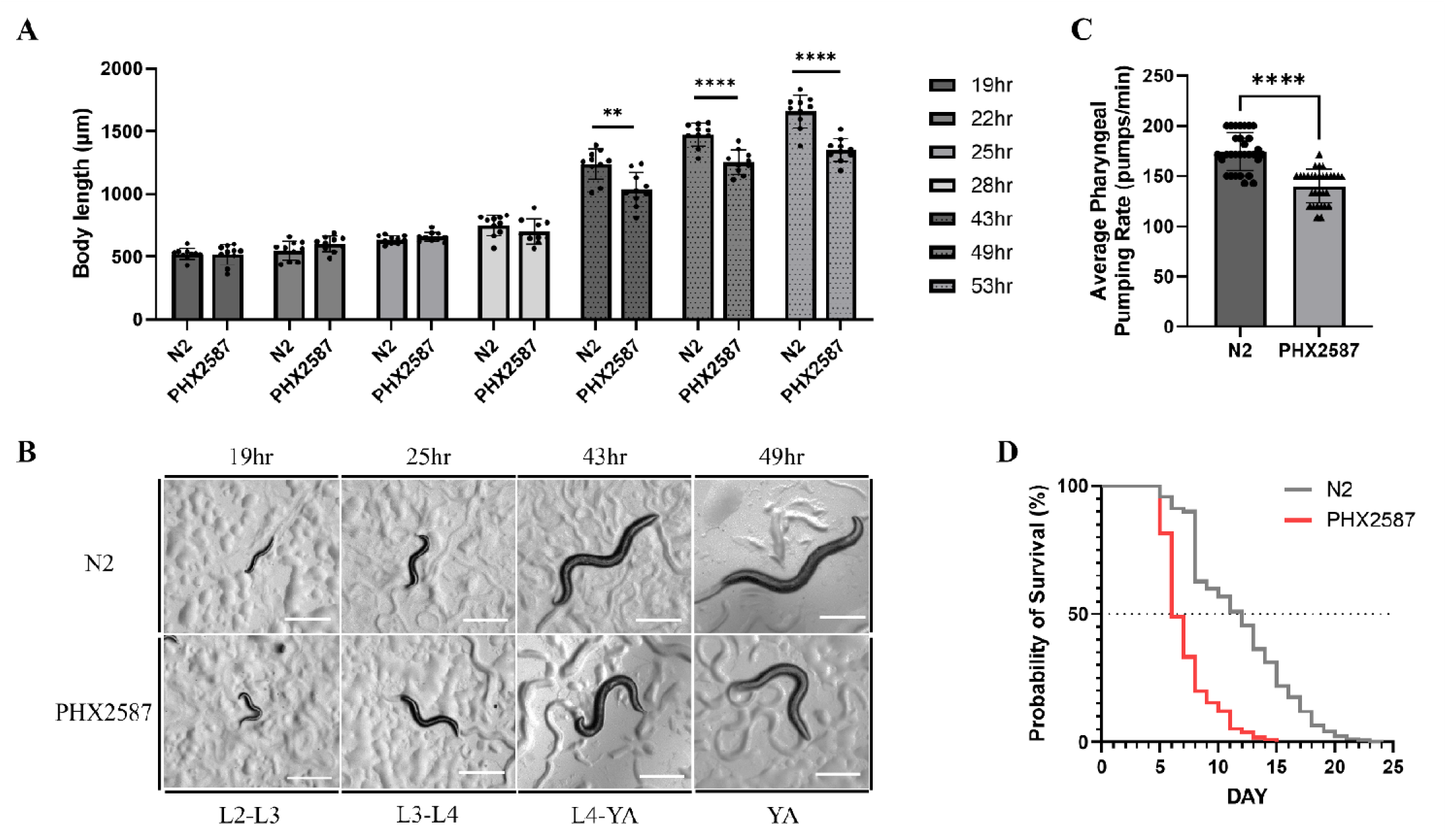
Morphological, physiological, and lifespan measurements in wac-deficient C. elegans. (A) Body length of N2 and *wac*-deficient PHX2587 worms measured at indicated time points (19–53 h) after L1 synchronization. Each dot represents one worm; bars indicate mean ± SD. Measurements were collected from three independent biological replicates, with 10 worms measured per genotype in each replicate (total n = 30 worms per genotype per time point). N2 and PHX2587 were compared at each time point using unpaired two– tailed t-tests. **p < 0.01; ****p < 0.0001. (B) Representative images of N2 and PHX2587 worms at matched developmental stages. Scale bars, 200 µm. (C) Pharyngeal pumping rate measured at 48 h after L1 synchronization (young adult stage) in N2 and PHX2587; bars indicate mean ± SD (n = 30). N2 and PHX2587 were compared using an unpaired two-tailed t-test. ****p < 0.0001. (D) Kaplan–Meier survival curves of N2 (n = 261) and PHX2587 (n = 313) worms. Survival differences were assessed using the log-rank (Mantel–Cox) test (p < 0.0001). Median survival was 12 days for N2 and 6 days for PHX2587.

### RNAi knockdown of cho-1 modifies body length and food-leaving behavior in PHX2587 worms

Because *cho-1* RNAi selectively reduced aldicarb-induced paralysis in PHX2587 worms, we next tested whether *cho-1* knockdown also modifies selected *wac*-associated phenotypes beyond aldicarb sensitivity. Body length was measured at 43 h and 50 h after L1 synchronization in worms grown on EV, *wac-1.2* RNAi, or *cho-1* RNAi plates. At both time points, *cho-1* RNAi had no significant effect on body length in N2 worms but significantly increased body length in PHX2587 worms (**Fig. 5A, B**). In contrast, the available *wac-1.2* RNAi clone did not significantly alter body length in N2 worms under these conditions, indicating that *wac-1.2* RNAi alone did not reproduce the PHX2587 growth-associated phenotype. We then assessed whether *cho-1* RNAi modifies food-leaving behavior. Following RNAi pretreatment, stage-matched young adult worms were transferred to OP50 assay lawns containing N2 progeny, with approximately 100–200 N2 eggs confirmed on each plate before the assay, and food-leaving behavior was quantified at 2 h and 24 h. At 24 h, PHX2587_EV worms showed reduced food-leaving behavior compared with N2_EV controls, whereas *cho-1* RNAi increased food-leaving behavior in PHX2587 worms while showing limited effects in N2 controls (**Fig. 5C**). Together with the aldicarb RNAi results, these findings support *cho-1* as a genotype-specific functional modifier of selected *wac*-associated cholinergic, growth-associated, and food-leaving phenotypes.

**Figure 5.**
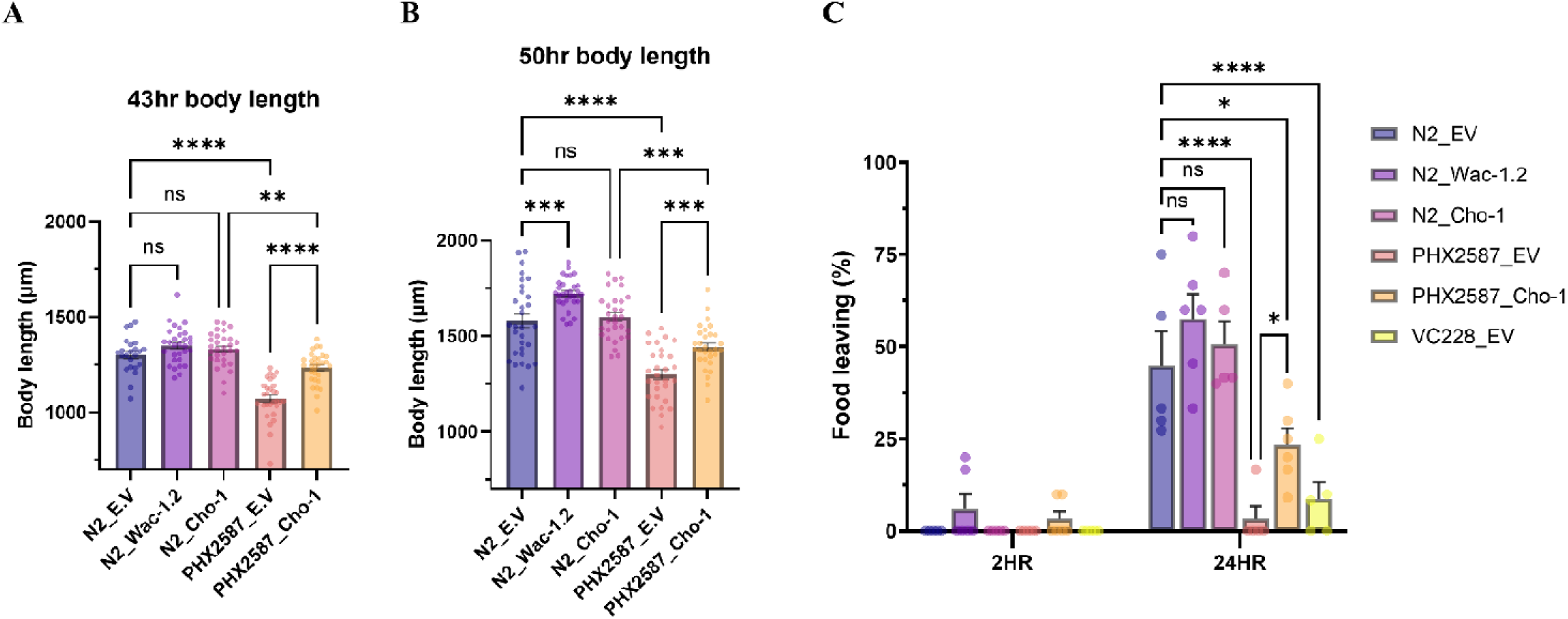
Effects of cho-1 RNAi on body length and food-leaving behavior in N2 and wac-deficient C. elegans. (A, B) Body length was measured at 43 h and 50 h after L1 synchronization in N2 and PHX2587 worms grown on empty vector control (E.V), *wac-1.2* RNAi, or *cho-1* RNAi plates. Body length data were analyzed by ordinary one-way ANOVA followed by Tukey’s multiple-comparisons test. Each dot represents an individual worm, and bars indicate mean ± SEM. Data were obtained from three independent biological replicates, with final analyzed sample sizes ranging from 23 to 30 worms per group. **p < 0.01; ***p < 0.001; ****p < 0.0001. (C) Food-leaving behavior was assessed in N2, PHX2587, and VC228 worms following E.V, *wac-1.2* RNAi, or *cho-1* RNAi pretreatment. Stage-matched young adult worms were transferred to OP50 assay lawns containing N2 progeny, with approximately 100–200 N2 eggs confirmed on each plate before scoring, and food-leaving behavior was quantified at 2 h and 24 h. Each dot represents one independent experiment, with 10 worms scored per experiment. Food-leaving data were obtained from 4–5 independent experiments per group. Food-leaving data were analyzed by two-way repeated-measures ANOVA followed by Tukey’s multiple-comparisons test. Bars indicate mean ± SEM. *p < 0.05; ****p < 0.0001.

## 4. Discussion

This study integrated behavioral, physiological, transcriptional, and functional analyses to define how loss of *wac* alters food-associated behavioral and cholinergic pathway function in *C. elegans*. *wac*-deficient PHX2587 worms exhibited a selective reduction in food-leaving behavior, while aggregation behavior was not detectably altered. This selective behavioral phenotype suggests that loss of *wac* may differentially affect specific cue-dependent behavioral responses, including pathways involved in exploratory or dispersal-related behavioral decisions. Because food-leaving behavior is influenced by neuromodulatory circuits, including dopaminergic and serotonergic pathways(Ishita et al., 2020; Sawin et al., 2000), the behavioral changes observed in *wac*-deficient worms may reflect altered neural regulation of behavioral state transitions. In parallel, *wac* deficiency led to reduced growth, decreased pharyngeal pumping, and shortened lifespan, reflecting systemic physiological impairment. Reduced pharyngeal pumping is particularly notable, as pharyngeal activity serves as a rhythmic neuromuscular readout of feeding physiology in *C. elegans* (Avery and You, 2012) and may reflect broader disruption of physiological regulation. Although pharyngeal pumping is not directly analogous to mammalian cardiovascular physiology, altered rhythmic physiological regulation in *C. elegans* may conceptually parallel autonomic dysregulation reported in ASD-associated conditions(Bujnakova et al., 2016; Cheng et al., 2020) Likewise, reduced growth and shortened lifespan in *wac*-deficient worms are consistent with broader developmental and physiological abnormalities reported in WAC-associated syndromes and related neurodevelopmental conditions, supporting a broader role for WAC in organismal homeostasis. Consistent with these findings, *Wac* loss-of-function in vertebrate models has been associated with developmental and behavioral abnormalities (Varvagiannis et al., 1993), suggesting a broader role for WAC in developmental regulation and organismal homeostasis. At the molecular level, stage-resolved transcriptional profiling revealed gene and stage-specific changes in cholinergic pathway components, with a pronounced coordinated shift emerging at the young adult stage. These stage-associated changes suggest that the effects of *wac* deficiency on cholinergic signaling may be developmentally regulated. Functional RNAi analysis further identified *cho-1* as a genotype-specific modifier of aldicarb sensitivity in PHX2587 worms. Extending this analysis, *cho-1* RNAi also increased body length and improved food-leaving behavior in PHX2587 worms, while having limited effects in N2 controls. Because *cho-1* encodes a high-affinity choline transporter required for acetylcholine synthesis(Matthies et al., 2006b), its genotype-specific effect suggests that altered presynaptic choline availability may contribute to altered cholinergic regulation observed in *wac*-deficient worms.

ASD (Manzo et al., 2025) and DESSH (Reynolds et al., 2024; Zhang et al., 2019) patients exhibit pronounced social and behavioral aberrations. Furthermore, there exists a significant overlap between DESSH and ASD. In a study conducted on four children with WAC variants, while all four exhibited developmental impairment and speech affliction, two were reportedly diagnosed with ASD (Alawadhi et al., 2021). *C. elegans* exhibit peculiar behavior where the adult worms avoid the food lawn crowded with progeny. This type of social behavior results from pheromonal cues generated by the progeny (Scott et al., 2017). Earlier studies utilizing the neuroligin/nlg-1 gene have identified depleted food leaving behavior, establishing it as a paradigm to test ASD linked behavior (Rawsthorne et al., 2021a). Food leaving assay to quantify social behavior in *C. elegans* and has been used to investigate genes linked to neurodevelopmental disorders, including ASD-associated genes, *nlg-1* (Rawsthorne et al., 2021a), *nrx-1*, and *chd-7* (Rawsthorne et al., 2021a; Rawsthorne et al., 2021b). Rawsthorne et al. showed that *nlg-1(ok259)* mutants exhibit reduced progeny-enhanced food leaving despite normal egg laying, normal progeny production, and no major defect in baseline locomotion, indicating that the phenotype dependent on complex intricate neuronal mechanisms. This phenomenon is similar in principle to human behavior that is impaired in ASD patients. Using progeny-preconditioned lawns and rescue experiments, they further demonstrated that the primary defect lies in the adult’s recognition and/or integration of progeny-derived social cues rather than impaired cue production by progeny (Rawsthorne et al., 2021a). Additional studies have further shown that larval-derived signals and pheromone-dependent cues contribute to the regulation of food– leaving behavior (Scott et al., 2017). In contrast, aggregation reflects a group-feeding state regulated by neuronal circuits that integrate sensory context and internal state (de Bono et al., 2002; Macosko et al., 2009b), suggesting that food leaving and aggregation represent distinct, though partially overlapping, behavioral outputs. The interpretation of food leaving behavior as a measure of social interaction is further supported by large-scale genetic screening studies in *C. elegans* (Rawsthorne et al., 2021b). A systematic analysis of ASD-associated gene orthologs demonstrated that progeny-induced food leaving reflects a complex, sensory-integrative behavior rather than simple locomotor output (Rawsthorne et al., 2021b). Mutations across diverse ASD– associated gene classes, including those involved in synaptic transmission, cell signaling, and epigenetic regulation, led to impaired food leaving behavior, while most mutants exhibited minimal defects in locomotion, feeding, or early development, further reinforcing that this assay selectively captures cue-dependent neural processing rather than general motor capacity. Consistent with these observations, the reduced food-leaving phenotype observed in *wac*-deficient worms suggests altered responsiveness to progeny-associated cues present on the food lawn. Although locomotor activity, sensory processing, and physiological state can influence food– leaving behavior, these factors are unlikely to fully account for the phenotype observed here because food leaving was assessed over an extended 24 h period, allowing sufficient time for worms to exit the lawn if the progeny-associated cue response was intact. In addition, aggregation on bacterial lawns was not detectably altered under our assay conditions, indicating that not all food-associated social behavioral outputs were affected by *wac* loss. This lack of an aggregation phenotype does not conflict with the reduced food-leaving response, because food-leaving and aggregation capture distinct behavioral outputs. Food-leaving reflects an adult response to progeny-associated cues that promotes leaving from the food lawn, whereas aggregation reflects group-feeding versus solitary behavior on bacterial lawns in the absence of progeny-derived cues. Thus, loss of *wac* appears to preferentially affect progeny-associated food-leaving behavior without broadly disrupting aggregation behavior under the conditions tested.

At the molecular level, WAC has been shown to regulate gene expression through its interaction with the RNF20/40 complex, which mediates histone H2B ubiquitination and transcriptional control (Zhang and Yu, 2011). In addition, WAC has been implicated in multiple cellular regulatory pathways, including mTOR signaling and autophagy, suggesting broader roles in cellular homeostasis and neuronal function (David– Morrison et al., 2016; Joachim et al., 2015). Together, these findings are consistent with a broader role for WAC in transcriptional regulation and cellular homeostasis, particularly in neuronal systems. However, the present data does not establish direct chromatin-level targeting of cholinergic pathway genes by WAC. Consistent with this regulatory role, *wac* deficiency resulted in coordinated changes in cholinergic pathway gene expression. Importantly, these changes were highly stage-associated. Early larval stages (L1–L3) showed modest or heterogeneous expression differences, whereas a pronounced and coordinated transcriptional shift emerged at the L4 and young adult stages. This temporal pattern suggests that the impact of *wac* loss becomes more prominent during later developmental stages associated with neuronal maturation. This stage-associated transcriptional shift is consistent with reports that WAC expression is developmentally regulated in the mammalian brain (Nishikawa et al., 2023), suggesting that disruption of WAC may have temporally restricted effects on neuronal gene regulation. In PHX2587, multiple cholinergic components exhibited coordinated expression changes at the young adult stage, whereas others showed gene-specific deviations during larval development, indicating both global and stage-associated regulatory effects. This dual pattern of regulation suggests that *wac* deficiency may influence both broad transcriptional programs and gene-specific regulatory mechanisms. Gene-level analysis indicates that this shift reflects coordinated transcriptional changes in cholinergic synaptic components. Presynaptic genes involved in acetylcholine synthesis, choline uptake, and vesicular transport, including *cha-1* (Alfonso et al., 1994), *cho-1* (Matthies et al., 2006a; Mullen et al., 2007), and *unc-17* (Alfonso et al., 1993), respectively, showed increased expression at later stages (L4–YA), suggesting elevated demand for neurotransmitter production. In parallel, postsynaptic receptor subunits and synaptic assembly factors, including *lev-1*, *lev-10*, *unc-29*, and *unc-38*, exhibited dynamic and stage-associated alteration, with pronounced upregulation at the young adult stage. The convergence of these presynaptic and postsynaptic changes supports the idea of cholinergic remodeling, potentially reflecting compensatory adaptation in response to *wac* deficiency. These coordinated changes across presynaptic and postsynaptic components suggest broader reorganization of cholinergic synaptic regulation rather than isolated gene-specific effects in *wac*-deficient worms.

Functional analysis using aldicarb assays confirmed that these transcriptional changes have physiological consequences. Aldicarb sensitivity is a well-established method for acetylcholine neurotransmission at the neuromuscular junction (Mahoney et al., 2006; Sammi et al., 2022). Consistent with the transcriptional remodeling observed in cholinergic pathway components, RNAi knockdown revealed that *cho-1* modulates aldicarb-induced paralysis in the *wac*-deficient background, identifying *cho-1* as a key genotype-specific functional modifier within the cholinergic pathway. This selective effect suggests a functional dependency that becomes evident in the context of *wac* deficiency. Because *cho-1* encodes a high-affinity choline transporter required for presynaptic acetylcholine synthesis (Matthies et al., 2006a; Mullen et al., 2007), this result suggests that presynaptic choline availability may become functionally more relevant in the *wac*– deficient background. This increased dependency is consistent with the upregulation of presynaptic cholinergic genes observed at later developmental stages, suggesting elevated demand for acetylcholine production. Furthermore, selective role of *cho-1* in PHX2587 worms suggests that altered synaptic acetylcholine availability may contribute to the observed phenotype. Consistent with previous reports, *cho-1* functions as a rate-limiting component of acetylcholine synthesis, and its disruption leads to activity-dependent cholinergic defects, while baseline phenotypes remain relatively mild due to compensatory choline synthesis pathways(Mullen et al., 2007). Taken together, these findings suggest that *wac* deficiency is associated with altered cholinergic regulation that is functionally modified by *cho-1*-mediated choline transport. Under these conditions, perturbation of *cho-1* may have a greater effect in the *wac*-deficient background than in N2 controls. This genotype-specific effect supports a functional relationship between *wac* deficiency, presynaptic choline transport, and selected cholinergic and behavioral phenotypes, without establishing *cho-1* as a direct transcriptional target of WAC. The extended *cho-1* RNAi analysis further supports this interpretation by linking *cho-1*-associated modulation to non-pharmacological phenotypes. In addition to reducing aldicarb sensitivity, *cho-1* RNAi increased body length and improved food-leaving behavior in PHX2587 worms, while having limited effects in N2 controls. These findings indicate that the functional consequence of *cho-1* perturbation is not restricted to aldicarb-induced paralysis but extends to growth-associated and cue-dependent behavioral phenotypes in the *wac*-deficient background. Importantly, although *cho-1* RNAi did not fully restore all phenotypes to the N2 control level, it produced consistent partial rescue across the outcomes tested, including aldicarb sensitivity, body length, and food-leaving behavior. Thus, *cho-1*-mediated choline transport appears to function as a genotype-specific modifier linking *wac* deficiency to altered cholinergic sensitivity, growth– associated progression, and progeny cue-dependent behavioral output.

Our previous work (Boonpraman et al., 2026) demonstrated that *wac* deficiency does not significantly alter total acetylcholine levels or acetylcholinesterase activity, but is associated with altered nicotinic acetylcholine receptor activity. The present study extends these prior observations by showing that *wac* loss is associated with stage-associated transcriptional changes in cholinergic pathway genes and by identifying *cho-1*– mediated choline transport as a genotype-specific modifier of aldicarb sensitivity, body length, and progeny– associated food-leaving behavior. In this context, the enhanced effect of *cho-1* knockdown in PHX2587 suggests that presynaptic choline uptake becomes a limiting factor under conditions of increased cholinergic demand. This interpretation is further supported by evidence that high-affinity choline transport constrains cholinergic output in systems with elevated synaptic activity (Bauche et al., 2016; Paolone et al., 2013; Parikh et al., 2013). In this model, postsynaptic receptor responsiveness may be altered, increasing the demand for acetylcholine signaling, while presynaptic mechanisms become limiting in their ability to sustain neurotransmitter supply. This creates a supply–demand imbalance in which synaptic output becomes increasingly dependent on the availability of choline and acetylcholine. Under these conditions, *cho-1* function may represent a genotype-sensitive point of vulnerability, consistent with its selective effects in the *wac*– deficient background. Together, these findings support a model in which *wac* deficiency is associated with compensatory-like changes in the cholinergic system, characterized by stage-associated transcriptional reorganization and increased functional dependence on presynaptic choline transport. The identification of *cho– 1* as a key functional node also highlights a broader principle of selective vulnerability within the cholinergic network. Although multiple pathway components exhibited transcriptional changes, only a subset exerted functional effects upon perturbation, suggesting that functional vulnerability may be concentrated at specific regulatory nodes rather than uniformly distributed across the pathway. This pattern is consistent with current models of neurodevelopmental disorders, in which disruption of broadly acting regulatory genes can produce pathway-specific functional vulnerabilities (De Rubeis et al., 2014; Satterstrom et al., 2020). The cholinergic system plays a central role in regulating attention, learning, and behavioral flexibility, and disruptions in cholinergic signaling have been implicated in ASD and related neurodevelopmental disorders (Deutsch et al., 2015). Nicotinic acetylcholine receptors are key modulators of synaptic transmission and circuit function, and their dysregulation can lead to altered neuronal responsiveness. In this context, our findings in *C. elegans* are consistent with a conserved role for cholinergic regulation in neural and behavioral function. Together with our prior observation of increased CHRNA7 expression in WAC-heterozygous mice(Boonpraman et al., 2026), these findings suggest that WAC may influence conserved cholinergic mechanisms across species, suggesting a connection between transcriptional regulation, synaptic function, and behavioral outcomes.

Finally, the stage-specific divergence in cholinergic gene expression suggests that the impact of *wac* deficiency is developmentally constrained rather than uniform. The pronounced transcriptional shift observed at the L4-to-adult transition indicates that WAC function may be particularly important during later stages of neuronal maturation. This developmental window coincides with periods of active neuronal maturation and synaptic remodeling in *C. elegans*. Disruption during this window may lead to persistent alterations in synaptic organization and function, even if earlier developmental stages exhibit only modest changes. While *C. elegans* does not fully recapitulate the complexity of ASD, its well-defined neural circuitry and conserved synaptic architecture provide a useful model system for identifying pathway-level mechanisms linking gene function to neural and behavioral outcomes (Cook et al., 2019a; Mizumoto et al., 2023; Rand, 2007; White et al., 1986b). Thus, insights gained from this model may provide broader insight into conserved principles of neural regulation and dysfunction.

Insufficient choline uptake and plasma levels have been reported in children diagnosed with ASD (Hamlin et al., 2013; Jennings and Basiri, 2022). qPCR studies of *cho-1* expression in worms indicate upregulation only in YA stage, suggesting a compensatory response that initiates at a later stage in the nematodes. Similarly, other components of cholinergic signaling exhibit dysregulation, reflecting a systemic disruption of neuronal components that may culminate in altered neurobehavioral implications in Autism. However, replication to humans regarding ASD manifestations is not absolute. This is likely due to differences between the two systems, as *C. elegans* has a simple nervous system and humans have a complex one, so the disease phenotype is not replicated in its entirety. On the other hand, our studies also validate *C. elegans* as an alternative model for research pertaining to human disorders.

Taken together, our results support WAC as a regulator of cholinergic-associated neural and behavioral phenotypes and suggest a functional relationship between stage-associated cholinergic transcriptional changes, *cho-1*-associated modulation, and food-leaving behavior. More broadly, this study suggests that disruption of chromatin-associated regulator may contribute to stage-specific reorganization of neurotransmitter-related pathways, potentially influencing selective behavioral phenotypes. While this study provides functional insight into cholinergic-associated changes in *C. elegans*, further studies in additional biological contexts and genes, including vertebrate models and human-relevant systems, will be important to evaluate the broader translational relevance of pathways influenced by WAC function.

## Acknowledgement

Strains were provided by the CGC, which is funded by NIH Office of Research Infrastructure Programs (P40 OD010440).

## Author credit statement

DK and NB performed experiments and compiled the MS, performing qPCR experiments. NK provided necessary support for the experiments. SRS conceived the idea and provided necessary inputs and edited the MS. In addition, SRS also performed RNAi studies.

## Data availability statement

The data supporting the findings of this study are available from the corresponding author upon reasonable request.

## Declaration of AI-assisted technology

The AI tool Grammarly was used to enhance the manuscript’s readability. The article has been thoroughly reviewed and verified by the authors. Authors take full responsibility for the interpretations and content presented.

## Figure Legends

**Supplementary Figure 1.**
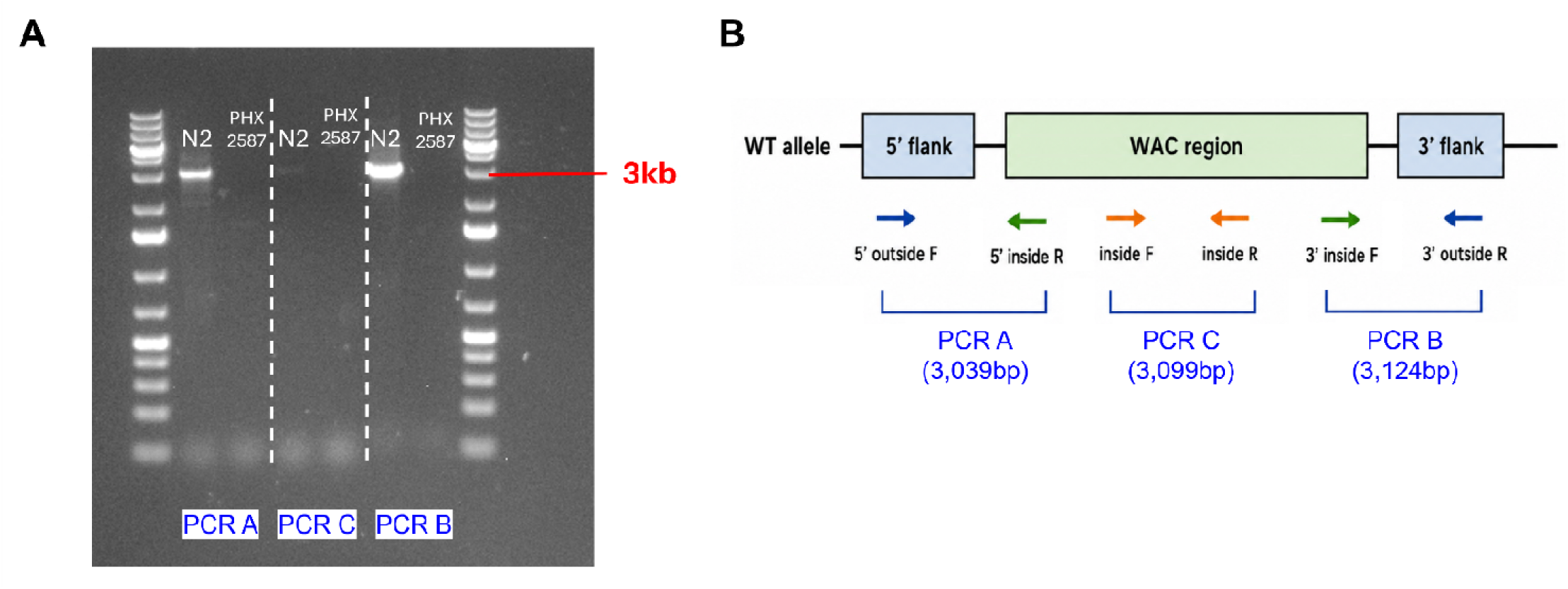
Genomic PCR validation of the PHX2587 wac deletion region. (A) Genomic PCR was performed to assess the presence of the corresponding wild-type *wac* genomic region in N2 and PHX2587 worms. WT-specific primer sets were used to amplify the 5′ flanking region (PCR A, 3,039 bp), an internal region within the wac locus (PCR C, 3,099 bp), and the 3′ flanking region (PCR B, 3,124 bp). PCR products of the expected size were detected in N2 genomic DNA, whereas corresponding WT-specific amplicons were not detected in PHX2587 genomic DNA. (B) Schematic representation of the WT *wac* genomic region and primer positions used for PCR validation. PCR A spans the 5′ outside forward and 5′ inside reverse primer pair, PCR C spans the internal forward and internal reverse primer pair, and PCR B spans the 3′ inside forward and 3′ outside reverse primer pair.

**Supplementary Figure 2.**
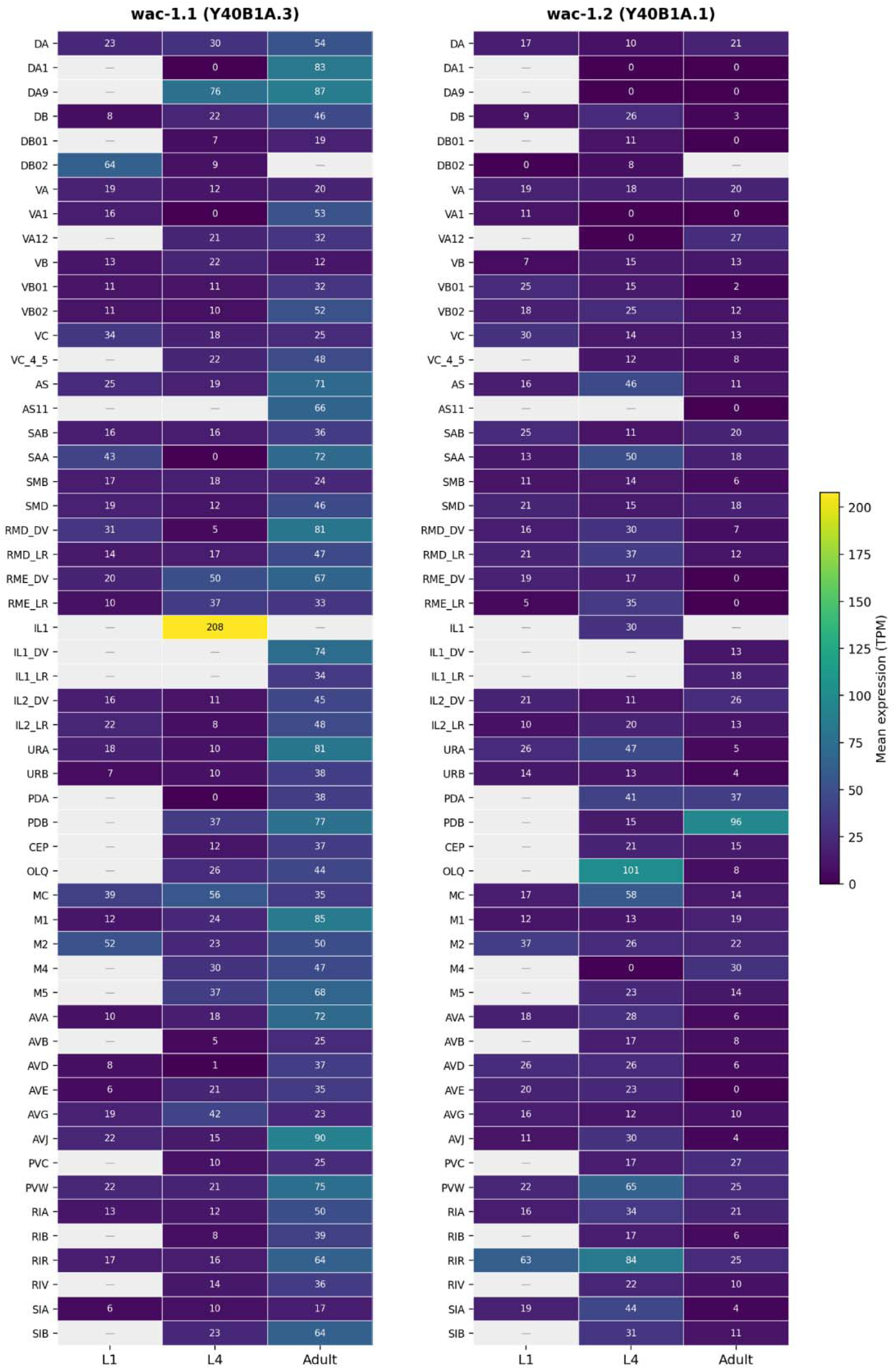
CeNGEN expression profiles of wac-1.1 and wac-1.2 across cholinergic neuron classes. Heatmap showing public CeNGEN single-cell RNA-seq expression profiles of *wac-1.1* and *wac-1.2* across cholinergic neuron classes at L1, L4, and adult hermaphrodite stages. *wac-1.1* was queried using the WormBase sequence name Y40B1A.3, and *wac-1.2* was queried using Y40B1A.1. Cholinergic neuron classes were defined based on acetylcholine neuron annotations. Color intensity represents the mean expression in TPM for each neuron class and developmental stage. Dashes indicate no detected expression or unavailable values in the queried dataset. These data indicate that *wac-1.1* and *wac-1.2* transcripts are detectable across multiple cholinergic neuron classes, supporting the plausibility that *wac* function may intersect with cholinergic circuits.

**Supplementary Table 1.** Primer sequence used for genomic PCR validation of the PHX2587 wac deletion region.

