## Supplementary Table 1 for "WAC loss alters food-associated behavior and physiological homeostasis with stage-associated cholinergic dysregulation in *Caenorhabditis elegans*"

| **Supplementary Table 1. Primer sequences used for genomic PCR validation of the PHX2587 *wac* deletion region.** | | | | |
| --- | --- | --- | --- | --- |
| **PCR assay** | **Target region** | **Primer name** | **Sequence (5′–3′)** | **Expected amplicon size** |
| PCR A | 5′ *wac* region | 5′ outside F | AGAGCTCTGGAACGAGAAAGAAAT | 3,039 bp |
| PCR A | 5′ *wac* region | 5′ inside R | TGGAAATTAGAGGGAAAAGCTCCA | 3,039 bp |
| PCR C | Internal *wac* region | Inside F | CCTTTTCCCATTGCTCAGCTTAAA | 3,099 bp |
| PCR C | Internal *wac* region | Inside R | AATGCCCTTTTAACAATTCTGGGG | 3,099 bp |
| PCR B | 3′ *wac* region | 3′ inside F | GCTCCATTGGCGCTATTATCTAGT | 3,124 bp |
| PCR B | 3′ *wac* region | 3′ outside R | AAAACGTGTAAAATCCCTTCCTCG | 3,124 bp |
